# Working-memory disruption by task-irrelevant talkers depends on degree of talker familiarity

**DOI:** 10.1101/372508

**Authors:** Jens Kreitewolf, Malte Wöstmann, Sarah Tune, Michael Plöchl, Jonas Obleser

**Affiliations:** Department of Psychology, University of Lübeck

**Author notes:** Correspondence should be addressed to:* Jens Kreitewolf and Jonas Obleser, Department of Psychology, University of Lübeck, Maria-Goeppert-Str. 9a, 23562 Lübeck, Germany.

**Keywords:** talker familiarity, working memory, irrelevant-speech task, attention, distraction

## Abstract

When listening, familiarity with an attended talker’s voice improves speech comprehension. Here, we instead investigated the effect of familiarity with a *distracting* talker. In an irrelevant-speech task, we assessed listeners’ working memory for the serial order of spoken digits when a task-irrelevant, distracting sentence was produced by either a familiar or an unfamiliar talker (with rare omissions of the task-irrelevant sentence). We tested two groups of listeners using the same experimental procedure. The first group were undergraduate psychology students (N=66) who had attended an introductory statistics course. Critically, each student had been taught by one of two course instructors, whose voices served as familiar and unfamiliar task-irrelevant talkers. The second group of listeners were family members and friends (N=20) who had known either one of the two talkers for more than ten years. Students, but not family members and friends, made *more* errors when the task-irrelevant talker was familiar versus unfamiliar. Interestingly, the effect of talker familiarity was not modulated by the presence of task-irrelevant speech: students experienced stronger working-memory disruption by a familiar talker irrespective of whether they heard a task-irrelevant sentence during memory retention or merely expected it. While previous work has shown that familiarity with an attended talker benefits speech comprehension, our findings indicate that familiarity with an ignored talker deteriorates working memory for target speech. The absence of this effect in family members and friends suggests that the degree of familiarity modulates memory disruption.

## Introduction

In natural situations, listeners are whelmed by a multitude of sounds that compete for attention. To comprehend a talker’s speech in the presence of competing distractors, both voice characteristics, such as the talker’s pitch, timbre, and articulatory style (reviewed by Diehl et al., 2004; Mathias and von Kriegstein, 2014), and listening goals, such as selective attention to and working memory of the talker’s speech (reviewed by Fritz et al., 2007; Shinn-Cunningham, 2008), guide the listener to focus on relevant sounds and to ignore irrelevant distractors.

One beneficial factor for speech comprehension under such conditions is familiarity with the talker’s voice. Several studies have shown that listeners are better at comprehending target speech when it is produced by a familiar compared to an unfamiliar talker (Holmes et al., 2018; Johnsrude et al., 2013; Kreitewolf et al., 2017; Levi et al., 2011; Newman and Evers, 2007; Nygaard and Pisoni, 1998, Souza et al., 2013). In the following, we refer to this phenomenon as the *familiarity benefit* (e.g., Johnsrude et al., 2013; Kreitewolf et al., 2017).

The familiarity benefit likely relies on listeners’ previous experience with the talker’s vocal characteristics which can be exploited to direct selective attention to target sounds in the auditory scene (Bressler et al., 2014; Kreitewolf et al., 2018). A conceivable implication of the familiarity benefit is that listeners do not only benefit from talker familiarity when their goal is to attend to the familiar talker’s speech but also when they want to *ignore* it. In other words, if the familiarity benefit is based on previous experience with the talker’s voice, then this experience might also help listeners to filter out distracting speech produced by this talker.

Only few studies have investigated whether talker familiarity helps listeners to ignore distracting, task-irrelevant speech. Johnsrude et al. (2013) presented listeners with two concurrent spoken sentences and asked them to report key words from the target sentence. Critically, the authors manipulated talker familiarity in these sentences such that either the attended (target), the unattended (masker), or none of the two was spoken by a highly familiar talker (i.e., the listeners’ spouses). Listeners correctly reported more key words from the target sentence when either the target or the masker sentence was spoken by a familiar compared to an unfamiliar talker, suggesting that talker familiarity facilitates both attending to target speech and ignoring distracting speech. In an earlier study, Newman and Evers (2007) asked listeners to attend to and immediately repeat (i.e., to shadow) speech from a target talker while a distracting talker was presented in the background. Listeners differed in their familiarity with one of the two concurrent talkers and whether or not they were told that they would hear a familiar voice (i.e., explicit vs implicit knowledge). In this study, talker familiarity was ensured by presenting speech produced by the listeners’ university professor. The results showed that listeners with explicit knowledge about talker familiarity made fewer shadowing errors than listeners who only had implicit knowledge or listeners who were not familiar with the talker at all. Yet, this benefit was limited to familiarity with the target talker. Unlike Johnsrude et al. (2013), listeners did not benefit from familiarity with the distracting background talker. Therefore, these two studies produced somewhat incongruent results with regard to the question of whether talker familiarity helps listeners to ignore distracting, task-irrelevant speech.

Here, we investigated the effect of talker familiarity on the distraction induced by task-irrelevant speech using a different, yet well-established experimental paradigm: the *irrelevant-speech task* (e.g., Colle and Welsh, 1976; Salamé and Baddeley, 1982). The irrelevant-speech task requires listeners to keep the serial order of to-be-attended target stimuli in working memory, while task-irrelevant, to-be-ignored speech is presented during memory retention. The number of incorrectly recalled targets is thought to increase proportionally to the distraction by task-irrelevant speech, making the irrelevant-speech task an effective paradigm to study memory disruption by distracting speech. To modulate talker familiarity, we used an adaptation of the irrelevant-speech task in which a task-irrelevant, to-be-ignored sentence was either spoken by a familiar or an unfamiliar talker (Fig. 1).

**Figure 1.**
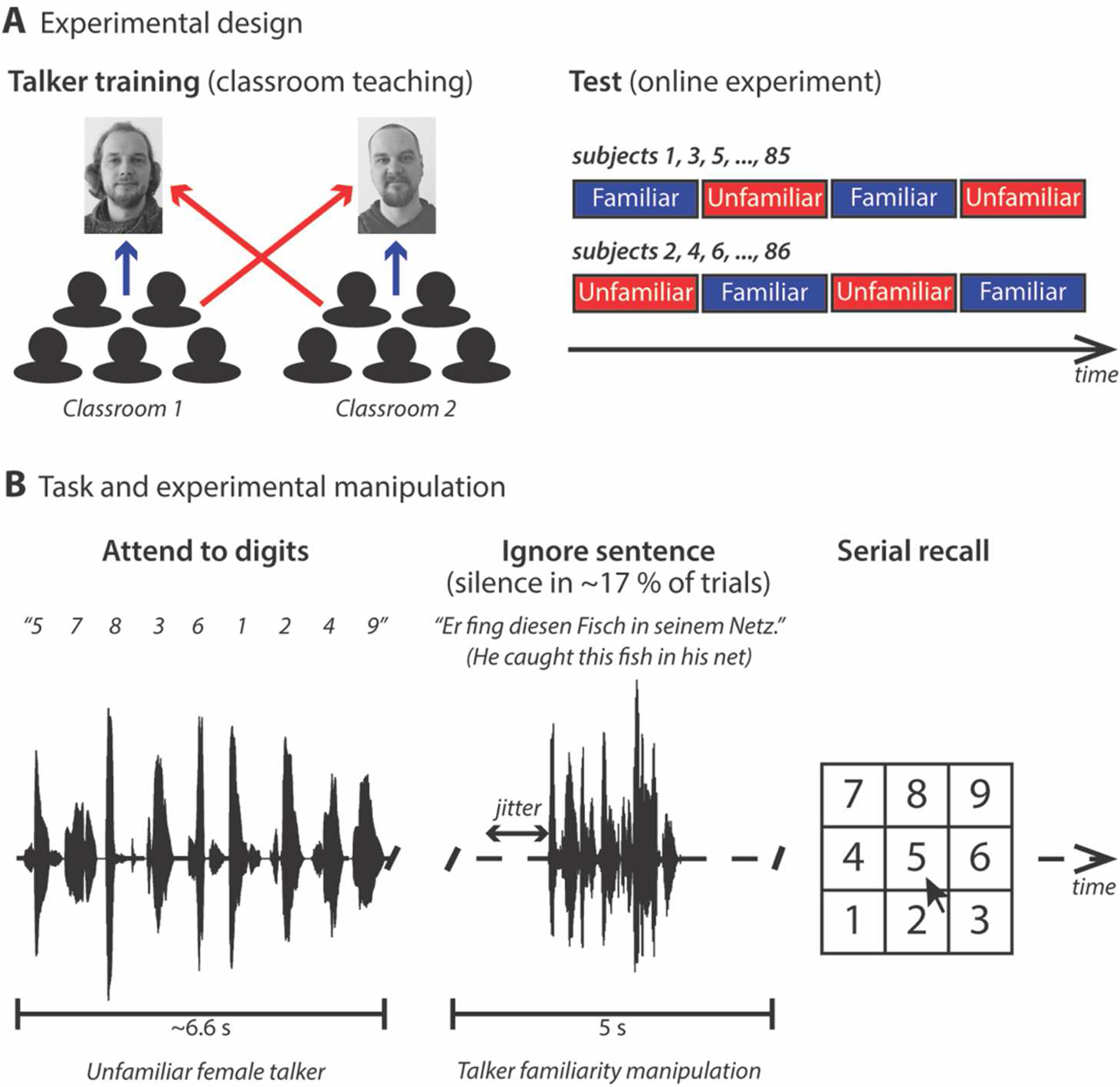
(**A**) Experimental design. Talker training in the students group was accomplished via classroom teaching. Students attended an introductory statistics course taught by one of two talkers. After the semester, the students as well as a group of family members and friends of the two talkers completed an online experiment. Familiar and unfamiliar talkers were presented in alternating blocks of trials. For half of the listeners, the experiment started with a familiar-talker block. (**B**) Task and experimental manipulation. In each trial of the online experiment, listeners heard the spoken digits 1 to 9 in random order followed by a task-irrelevant sentence. The onset of the sentence was jittered. In about 17 % of the trials, the sentence was omitted. The listeners’ task was to select the digits in the order of presentation from a visually presented number pad. The same task-irrelevant sentences were spoken by both talkers. Digits were spoken by an unfamiliar female talker.

The major objective of this study was to test whether familiarity with a task-irrelevant talker affects the serial recall of attended target speech. One possibility is that talker familiarity helps listeners to filter out irrelevant speech. This should manifest in *fewer* recall errors when the task-irrelevant talker is familiar versus unfamiliar. Such finding would not only be in line with Johnsrude et al. (2013) but also consistent with the idea of proactive filtering: when a distractor is known or anticipated, the filter can be applied even before the distractor appears (e.g., Noonan et al., 2016; Ruff and Driver, 2006). Interestingly, proactive filtering is beneficial when a distractor is present, but behaviorally costly when an expected distractor is omitted (Marini et al., 2013).

To test the possibility that listeners proactively filter out irrelevant speech produced by the familiar talker, we blocked the presentation of familiar and unfamiliar talkers and omitted the task-irrelevant sentence on rare occasions. If listeners proactively filtered irrelevant speech from a familiar talker, we would observe fewer recall errors on trials where the task-irrelevant sentence is spoken by a familiar versus unfamiliar talker (familiarity benefit) and more recall errors on trials where a familiar versus unfamiliar distracting talker is anticipated but the irrelevant sentence is omitted (familiarity deficit).

An alternative possibility is that task-irrelevant speech produced by a familiar talker captures attention and draws it away from items in memory. Previous work on the irrelevant-speech effect has shown that distractors of high familiarity, such as the listener’s own name (Röer et al., 2013) and the listener’s native language (Ellermeier et al., 2015), enhance memory disruption. Possibly, familiar distractors constitute salient stimuli that involuntarily draw attention resources away from the serial memory of target items. If this is also the case for familiar voices, we would observe *more* recall errors when the task-irrelevant sentence is spoken by a familiar versus unfamiliar talker; however, we would expect no effect of talker familiarity on trials where the task-irrelevant sentence is omitted.

A third possibility is that the effect of talker familiarity is not modulated by the actual presentation of a task-irrelevant sentence. That is, talker familiarity affects working memory irrespective of whether the familiar talker’s distracting speech is heard or merely expected. Such finding would be difficult to explain by (proactive) filtering or attentional capture of the familiar talker’s speech; instead, it would rather speak to more general differences in how familiar and unfamiliar task-irrelevant talkers affect working memory of target speech.

Based on the collective results from previous studies, one might draw the conclusion that the degree of familiarity plays an important role for the distraction by irrelevant speech and that listeners only benefit from high (Johnsrude et al., 2013) but not moderate familiarity (Newman and Evers, 2007) with a distracting talker. Yet, these studies are difficult to compare since they differ markedly in their experimental procedures, including stimuli and task instructions.

Here, we tested the effect of the degree of familiarity by comparing two groups of listeners that performed the exact same irrelevant-speech task. Importantly, these two groups differed in their degree of familiarity with similar magnitudes of familiarity as in previous studies. Specifically, we compared a group of students who heard irrelevant speech produced by one of their course instructors (i.e., moderate familiarity similar to Newman and Evers, 2007) with a group of the course instructors’ family members and friends (i.e., high familiarity similar to Johnsrude et al., 2013).

## Methods

### Participants

Two groups of listeners participated in this study. The first group were *N* = 66 undergraduate psychology students (59 females, mean age 23.11 yrs, age range 17-48 yrs; see Table I for details) who had received classroom instructions by one of two talkers. Classroom teaching comprised a total of fourteen 90-minute sessions (of which all participants attended at least nine; see Table I). The second group of listeners were *N* = 20 family members and close friends of either one of the two talkers (8 females, mean age 39 yrs, age range 30-65 yrs; see Table I for details), who did not receive classroom instructions. All listeners were native German speakers. Students gained course credit for their participation; family members and friends were paid € 10. The experimental procedures were approved by the ethics committee of the University of Lübeck.

**Table I.** Details of listener groups.

1: Students
| *familiar talker* | *N* | *gender*<br>(female/male) | *mean age*<br>(yrs) | *age range*<br>(yrs) | *mean familiarity*<br>(sessions) | *familiarity range*<br>(sessions) |
| --- | --- | --- | --- | --- | --- | --- |
| **JK** | 32 | 28/4 | 24.27 | 17-48 | 13.19 | 9-14 |
| **MP** | 34 | 31/3 | 22.09 | 18-30 | 12.94 | 10-14 |

**Group 2: Family and friends**
| *familiar talker* | *N* | *gender*<br>(female/male) | *mean age*<br>(yrs) | *age range*<br>(yrs) | *mean familiarity*<br>(yrs) | *familiarity range*<br>(yrs) |
| --- | --- | --- | --- | --- | --- | --- |
| **JK** | 10 | 3/7 | 39.10 | 30-65 | 23.10 | 14-34 |
| **MP** | 10 | 5/5 | 38.90 | 32-42 | 25.30 | 11-40 |

### Stimuli

The to-be-attended speech stimuli were recordings of the German digits 1 to 9, which we had used in previous studies (Obleser et al., 2012; Wöstmann and Obleser, 2016; Wöstmann et al., 2017). All digits were spoken by a native German female talker (mean fundamental frequency, *f0*, 192 Hz). None of the listeners was familiar with the talker’s voice. Digits had an average duration of 0.6 s (ranging from 0.5 to 0.7 s) and were concatenated with an onset-to-onset delay of 0.75 s. The resulting digit streams had an average duration of 6.6 s.

For the task-irrelevant speech, we used a German version of the speech-in-noise sentences (Erb et al., 2012) adopted from Kalikow et al. (1977). The same 50 sentences were recorded from two male talkers who were the authors JK and MP (both native German speakers). The mean *f0*, averaged across all sentences, was 93 Hz for talker JK, and 85 Hz for talker MP. Sentences produced by JK had an average duration of 2.17 s (ranging from 1.83 to 2.58 s); sentences produced by MP had an average duration of 2.19 s (ranging from 1.91 to 2.72 s). All sentences and digit streams were normalised to the same root mean square (RMS) dB full-scale amplitude using MATLAB (version 8.6, MathWorks, United States). On a given trial, the onset of the task-irrelevant sentence was delayed by 1409 ms (± 400 ms) so that, on average, sentences were centered in the middle of a 5-second memory retention phase (Fig. 1B).

### Procedure

The listeners performed an online experiment implemented in Labvanced (Scicovery GmbH, Osnabrück, Germany) that used an adaptation of the irrelevant-speech paradigm (e.g., Colle and Welsh, 1976; Jones and Morris, 1992). The online experiment was executed in the browser in full-screen mode. Online experiments allow for the rapid collection of large datasets (e.g., Buhrmester et al., 2011) and have been shown to produce reliable data in several areas of behavioral research, including linguistics (e.g., Saunders et al., 2013) and psychoacoustics (e.g., McPherson and McDermott, 2018). Here, the online experiment had the additional advantage to prevent direct contact between the listeners and one of the two task-irrelevant talkers immediately before the start of the experiment, which would have otherwise contaminated our manipulation of talker familiarity.

All listeners completed the experiment within one hour. On each trial, listeners heard the nine spoken digits in random order followed by a task-irrelevant sentence (Fig. 1B), while a fixation cross was presented on the computer screen. In about 17 % of trials (i.e., 20 out of 120 trials), silence was presented instead of task-irrelevant speech. Five seconds after the offset of the digit stream (i.e., at the end of the memory retention phase), a number pad consisting of the digits 1 to 9 was visually presented. Listeners were asked to select the digits in the order of their presentation. Each visually presented digit disappeared directly after it had been selected. This was done to avoid that the same digit could be selected more than once per trial. After the selection of the ninth digit, the next trial started with a delay of 500 ms. No feedback was given.

Listeners were asked to perform the online experiment in a quiet setting, to use a computer (no tablets, smart phones, etc.), and to listen to the sounds using headphones. Prior to the start of the experiment, listeners were instructed to silently rehearse the digit stream during the memory retention phase, to keep their eyes open and not to speak the digits aloud during a trial. Listeners could adjust the loudness of the sounds to a comfortable level. They were asked not to change the sound level during the experiment.

The experiment comprised four blocks (Fig. 1A, ‘Test’). Each block consisted of 30 trials; 25 trials with a distracting, task-irrelevant sentence (distractor trials) and 5 trials with silence in the memory retention phase (no-distractor trials). In each block, no-distractor trials were pseudo-randomly interspersed with distractor trials with the restriction that the first no-distractor trial within a block could not occur within the first five trials and that two no-distractor trials could not occur in succession. The task-irrelevant familiar and unfamiliar talkers were presented in alternating blocks of trials. Talker familiarity was not made explicit (i.e., listeners were not told that they would hear a familiar talker’s voice). Half of the listeners started with a familiar-talker block; the other half started with an unfamiliar-talker block (Fig. 1A, ‘Test’). Note that no-distractor trials were acoustically identical for the task-irrelevant familiar and unfamiliar talker. However, the blocked presentation of the familiar and the unfamiliar talker allowed us to test whether the *infrequent* presentation of silence in the memory retention phase would affect performance differently when a familiar compared to an unfamiliar talker’s voice was expected.

In the first and second half of the experiment (consisting of one familiar-and one unfamiliar-talker block), the same combinations of digit streams and task-irrelevant sentences were presented (but each task-irrelevant sentence was spoken by one talker in the first half and by the other talker in the second half of the experiment). This was done to ensure that differences in performance between blocks were due to familiarity with the task-irrelevant talker and not due to differences in the memorability of digit streams or the distractibility of task-irrelevant sentences. To reduce item-specific learning, we ensured that the trial order was always different in familiar-and unfamiliar-talker blocks.

### Statistical analyses

To assess listeners’ memory of the serial order of digits, we considered digits recalled at their respective position of presentation as *correct* and all other responses as *incorrect*. As a measure of distraction by task-irrelevant speech, we counted the number of errors per trial (0-9). All statistical analyses were carried out in R (R Core Team, 2017) using RStudio (version 1.1.383).

To overcome problems related to the unequal number of trials with and without a distractor in the memory retention phase, we fitted generalized linear mixed-effects models as implemented in the *lme4* package (Bates et al., 2015) to the number of errors per trial using Poisson regression (log link function; treating the number of errors as count data).

We followed an iterative model fitting procedure: starting with a minimal model that only included the random intercepts for subjects, we first added fixed-effects and then random-effects terms in a stepwise fashion. Fixed-effects terms were added in the order of their conceptual importance (i.e., Talker Familiarity, Distractor, Listener Group, and interactions between these factors; see below). Random-effects terms included the random intercepts for sentences and the subject-and sentence-specific random slopes for all significant main factors and interactions. After each step, we fitted the model using maximum-likelihood estimation, and assessed the change in model fit using likelihood-ratio tests. Model terms that significantly improved the model fit were kept in the model, non-significant terms were dropped (unless they were involved in higher-order interactions), resulting in a best-fitting model.

To investigate the potential effects of *Talker Familiarity* (familiar vs unfamiliar task-irrelevant talker), *Distractor* (distractor vs no-distractor trial), and *Listener Group* (students vs family and friends) on the number of errors, we modeled these predictors as fixed effects using deviation coding. To explore significant interaction terms, we performed post-hoc comparisons using Tukey’s range tests as implemented in the *lsmeans* package (Lenth, 2016). We report unstandardized coefficients *b* to provide an estimate of effect size for fixed effects. Note that Poisson regression operates on a log transform of the dependent measure. Coefficients are therefore given in log-scale units. For significant random-effects terms, we report the likelihood-ratio test comparing the more complex model that includes the random-effects term with the simpler model excluding the term.

To enhance the interpretability of non-significant effects in particular, we calculated Bayes Factors (BFs) using the *brms* package (Bürkner, 2016). When comparing two statistical models, the BF indicates how many times more likely the observed data are under the more complex model (including a particular model term of interest) compared to the simpler model (excluding the model term of interest). In accordance with Jeffreys (1939/1961), a BF of 0.33 or smaller is interpreted as providing evidence in favor of the null hypothesis, while a BF of 3 or larger is interpreted as evidence against it.

## Results

Figure 2 (A) shows the average proportions of errors as a function of digit position for distractor and no-distractor trials. Several basic observations can be made based on the data shown in this figure. Descriptively, listeners made fewer errors for digits presented at initial and final positions which is likely due to primacy and recency effects (e.g., Jones and Macken, 1993; Röer et al., 2013; Salamé and Baddeley, 1982; Schlittmeier et al., 2011; Wöstmann and Obleser, 2016). Interestingly, differences between distractor and no-distractor trials were most pronounced for digits presented in the second half of the digit stream (i.e., between the fifth and eighth digit), and digits presented at these positions were generally more difficult to recall. Another observation is that, averaged across digit positions, listeners made more errors in distractor compared to no-distractor trials (Fig. 2B).

**Figure 2.**
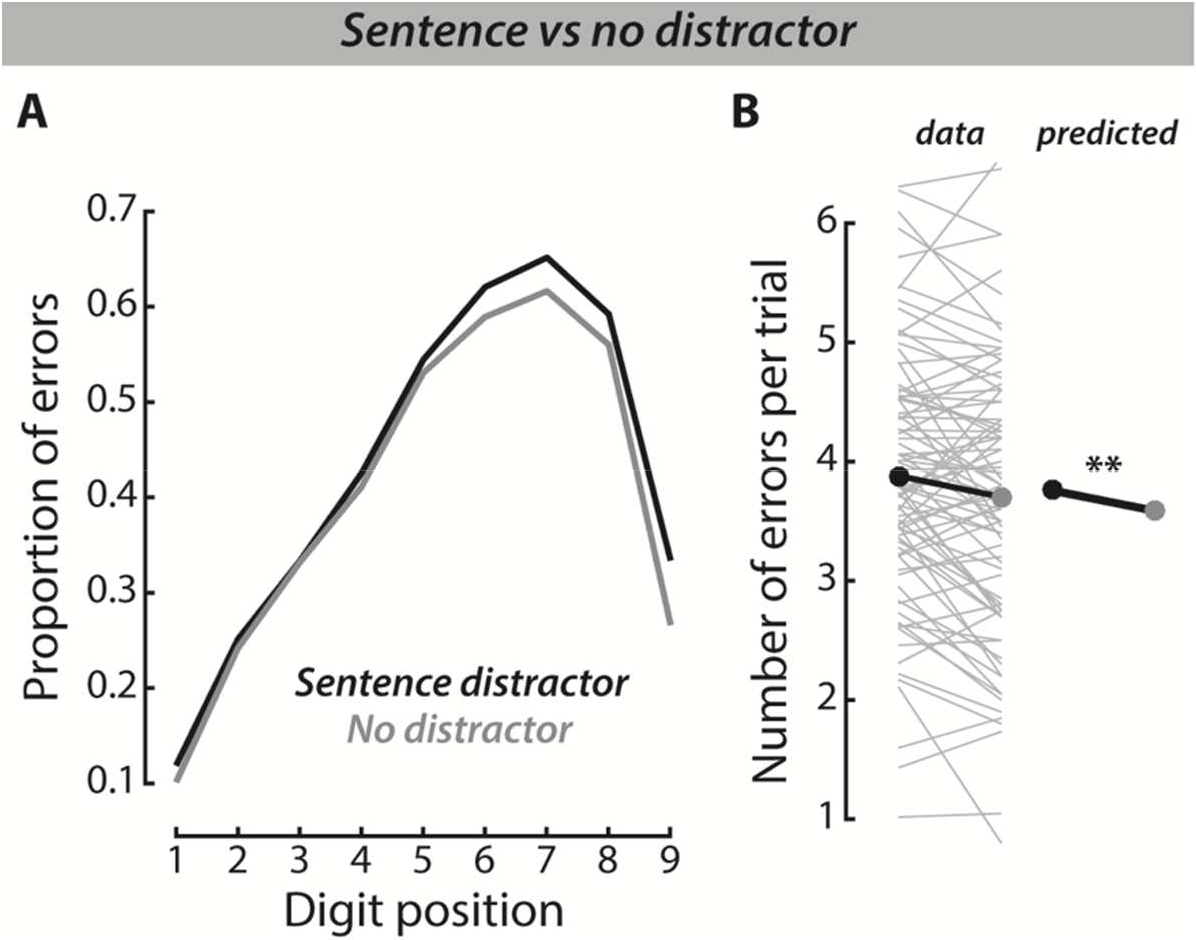
Distraction by task-irrelevant speech. (**A**) Average proportions of errors in sentence-distractor (black) and no-distractor trials (gray) as a function of digit position. (**B**) Left: number of errors per trial averaged across digit positions for sentence-distractor (left; black) and no-distractor trials (right; gray). Individual listeners’ data are shown as gray lines. Across-listener averages are shown by dots and are connected by the thick black line. Right: number of errors per trial as predicted by the best-fitting model (i.e., controlling for the effects of all other model terms; see Table II for a summary of model terms). Significant effects are denoted by asterisks: ** *p* < 0.01.

### Distraction by task-irrelevant speech

To test the effect of Distractor, amongst other things, on the number of errors per trial, we fitted linear mixed-effects models. In sum, the best-fitting model included the three main factors Distractor, Talker Familiarity, and Listener Group as well as the interaction between the factors Talker Familiarity and Listener Group as fixed-effects terms (see Table II for a summary of fixed-effects terms). The random-effects terms included the random intercepts for subjects and sentences as well as the subject-specific random slopes for Talker Familiarity.

**Table II.** Fixed-effects terms included in the best-fitting model.

| ***Fixed effect*** | ***Estimate b*** | ***Standard error SE*** | ***p-value*** |
| --- | --- | --- | --- |
| (Intercept) | 1.30057 | 0.03974 | < 2e-16 |
| Talker Familiarity (familiar) | -0.04718 | 0.02383 | 0.04772 |
| Listener Group (students) | -0.01155 | 0.07803 | 0.88236 |
| Distractor (distractor trial) | -0.04704 | 0.01676 | 0.00501 |
| Talker Familiarity × Listener Group | 0.09499 | 0.04737 | 0.04491 |

Note that the results are in log-scale units. Reference condition is given in brackets.

The inclusion of Distractor in the best-fitting model (*Z* = −2.81; *p* = 0.005; *b* = −0.047; *BF* = 7.6) demonstrated the *irrelevant-speech effect:* presentation of a task-irrelevant sentence within the memory retention phase was indeed more distracting than a silence period. The sentence-specific random intercepts were also included in the best-fitting model (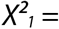; *p* = 0.002; *BF* = 4.09), suggesting that the task-irrelevant sentences differed in distractibility.

### Moderate but not high talker familiarity disrupts memory of target speech

The main aim of the present study was to investigate the effect of familiarity with a task-irrelevant talker on the serial recall of target digits. Figure 3 shows how the average proportion of errors evolves over digit positions in familiar-and unfamiliar-talker blocks (Fig. 3A) and the number of errors per trial averaged across digit positions for task-irrelevant familiar and unfamiliar talkers (Fig. 3B).

**Figure 3.**
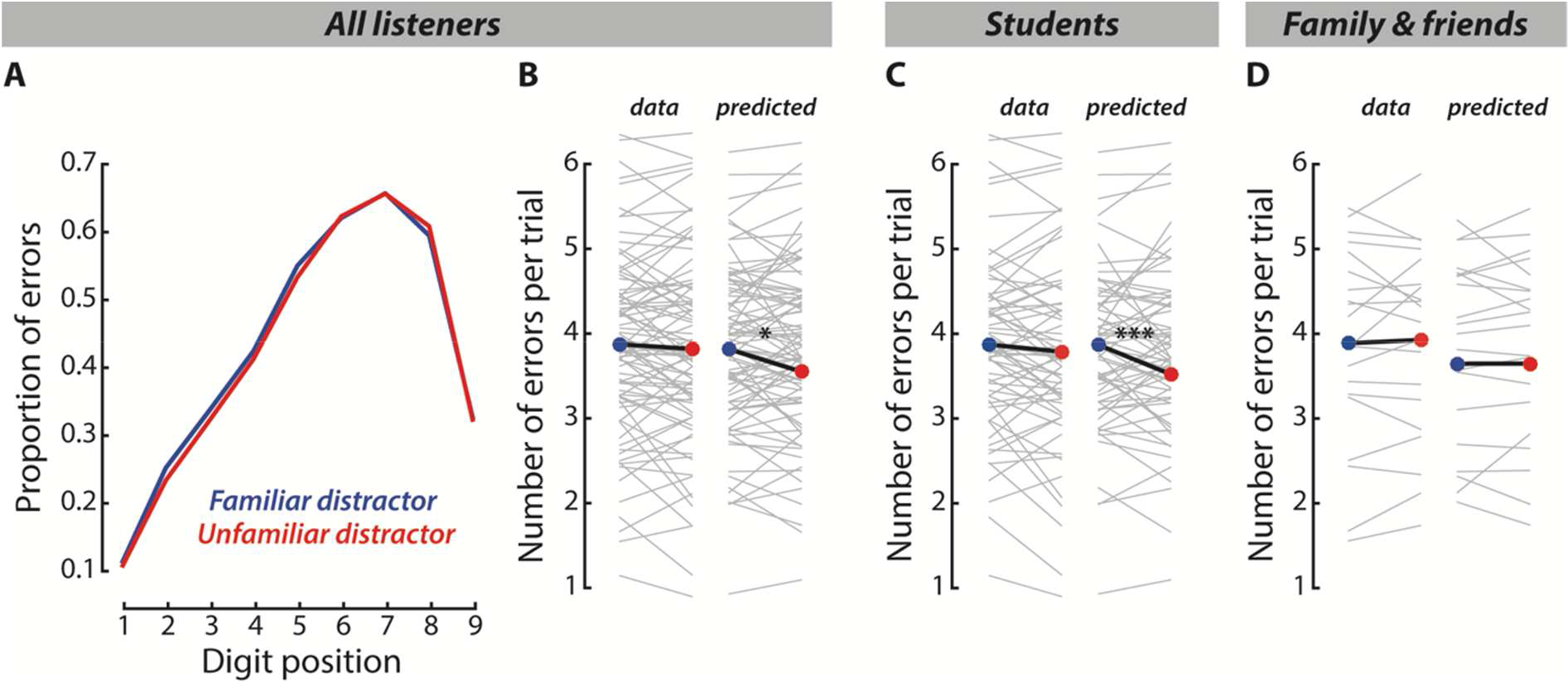
Effect of talker familiarity in all listeners (A, B) as well as in students (C) and in family members and friends (D). (**A**) Average proportions of errors in familiar-(blue) and unfamiliar-talker blocks (red) as a function of digit position. (**B, C, D**) Left: number of errors per trial averaged across digit positions for familiar-(left; blue) and unfamiliar-talker blocks (right; red). Individual listeners’ data are shown as gray lines. Across-listener averages are shown by colored dots and are connected by the thick black line. Right: number of errors per trial as predicted by the best-fitting model. Gray lines show the pairs between the random intercepts for subjects and the subject-specific random slopes for talker familiarity. Significant effects are denoted by asterisks: * *p* < 0.05; *** *p* < 0.001.

The best-fitting model included the factor Talker Familiarity (*Z* = −1.98; *p* = 0.048; *b* = −0.0472; *BF* = 1.58) and the interaction between the factors Talker Familiarity and Listener Group (*Z* = 2.01; *p* = 0.045; *BF* = 0.58). Post-hoc comparisons revealed that there was a significant effect of talker familiarity in the students group (*Z* = −4.07; *p* < 0.001; *b* = −0.0947; *BF* = 13.71) but not in the group of family members and friends (*Z* = 0.01; *p* = 0.994; *b* = 0.0003; *BF* = 0.04). This means that students (Fig. 3C) but not family members and friends (Fig. 3D) made *more* errors when they were familiar with the task-irrelevant talker than when they were not.

Notably, the sample size was much smaller in the group of family and friends (*N* = 20) than in the student group (*N* = 66). However, the small Bayes Factor of 0.04 provides strong evidence for the absence of a familiarity effect for family and friends. It is therefore unlikely that the comparably small sample was responsible for the lack of a familiarity effect in this group of listeners.

The best-fitting model also included the subject-specific random slopes for Talker Familiarity (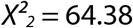; *p* < 0.001; *BF* = 17.72), suggesting that the effect of talker familiarity differed across listeners. Interestingly, however, the best-fitting model did not include the interaction between the factors Talker Familiarity and Distractor: compared to the simpler model, the inclusion of the fixed-effects term for the Talker Familiarity-by-Distractor interaction did not significantly improve the model fit (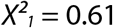; *p* = 0.434; *BF* < 0.001). Thus, the effect of talker familiarity (higher distraction in familiar-than unfamiliar-talker blocks) was not modulated by the presentation of a task-irrelevant sentence within the memory retention phase. This finding suggests that the mere expectation of a (moderately) familiar talker in the memory retention phase suffices to disrupt listeners’ working memory of target speech.

### Familiarity effects are talker-specific

By design, it is possible that any familiarity effect would only be driven by one of our two task-irrelevant talkers. To investigate this possibility, we carried out a control analysis in which we added the factor Familiar Talker (JK vs MP) and, critically, the interaction between Familiar Talker and Talker Familiarity as a fixed-effects terms to the best-fitting model. The inclusion of Familiar Talker did not significantly improve the model fit compared to the best-fitting model (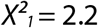; *p* = 0.138; *BF* = 0.32), but the inclusion of the interaction term between Familiar Talker and Talker Familiarity did (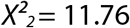; *p* = 0.003; *BF* = 5.33). Post-hoc comparisons revealed that listeners who were familiar with talker MP showed a detrimental effect of talker familiarity on their serial recall of target digits (*Z* = −3.67; *p* < 0.001; *b* = −0.1057; *BF* = 13.61). This was not the case for listeners who were familiar with talker JK (*Z* = 0.44; *p* = 0.663; *b* = 0.013; *BF* = 0.03).

While these results suggest that memory disruption by a familiar talker might be talker-specific, they cannot explain our main finding of stronger memory disruption by moderate (but not high) familiarity with the task-irrelevant talker. First, the factor Familiar Talker did not modulate the Talker Familiarity-by-Listener Group interaction: the inclusion of the three-way interaction between Familiar Talker, Talker Familiarity, and Listener Group did not significantly improve the model fit compared to the simpler model 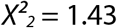; *p* = 0.49; *BF* = 0.04). Second, the interaction between Talker Familiarity and Listener Group remained a significant predictor for the number of errors per trial even after inclusion of the Familiar Talker-by-Talker Familiarity interaction (*Z* = 2.07; *p* = 0.038) with higher memory disruption by the familiar talker in the student group (*Z* = −4.20; *p* < 0.001; *b* = −0.0926; *BF* = 27.61) but not in the group of family members and friends (*Z* = −0.001; *p* = 0.999; *b* = −4.49e-5; *BF* = 0.04). Third, and most importantly, the identity of the familiar talker was balanced across listeners; that is, a similar number of listeners was familiar with either one of the two talkers (see Table I for details). Our experimental design therefore inherently controlled for any talker-specific effects on the main effect of talker familiarity as well as the Talker Familiarity-by-Listener Group interaction.

### No effect of listener group and block order

In total, students did not differ from family members and friends in their serial recall of digits. Compared to the simpler model, the inclusion of the factor Listener Group did not significantly improve the model fit (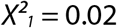; *p* = 0.888; *BF* = 1.11). Note, however, that the BF shows no evidence for either the absence or presence of the effect of Listener Group. It is thus possible that the non-significant difference between the two listener groups was due to the rather small sample of family members and friends.

To investigate potential effects of the presentation order of familiar and unfamiliar talkers (Fig 1A, ‘Test’), we carried out a second control analysis in which we added the factor *Block Order* (familiar first vs unfamiliar first) as a fixed-effects term to the best-fitting model. Compared to the respective simpler model, neither the main effect of Block Order (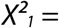 0.04; *p* = 0.843; *BF* = 0.06) nor the interaction term between the factors Block Order and Talker Familiarity (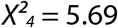; *p* = 0.223; *BF* = 0.05) significantly improved the model fit. Thus, neither the overall performance nor the effect of talker familiarity depended on whether listeners started the experiment with a familiar-or an unfamiliar-talker block.

## Discussion

The present study used a variant of the irrelevant-speech task to investigate the effect of familiarity with a distracting, task-irrelevant talker on the serial recall of target speech. The main finding was that listeners made *more* recall errors in blocks of trials with a familiar compared to an unfamiliar distracting talker. Critically, this effect depends on the degree to which listeners are familiar with this task-irrelevant voice: only listeners with moderate (i.e., students) but not high familiarity (i.e., family and friends) showed stronger working-memory disruption by talker familiarity. Interestingly, however, the effect of talker familiarity does not depend on the presence of task-irrelevant speech during memory retention: students experienced stronger working-memory disruption irrespective of whether they heard a task-irrelevant sentence produced by the familiar talker (in most of the trials) or merely expected it (in a small subset of trials).

### Familiarity with a distracting talker improves comprehension, but disrupts working memory of target speech

Two previous studies had investigated the distraction by familiar talkers’ speech. One study (Newman and Evers, 2007) found no benefit from moderate familiarity with a distracting background talker (i.e., the listeners’ university professor) when listeners had to shadow target speech. Another study (Johnsrude et al., 2013), however, showed that listeners’ comprehension of target speech does benefit from high familiarity with a distracting talker (i.e., the listener’s spouse)—a finding that has been recently extended to familiarity with accented speech (Senior and Babel, 2018). Together, these studies suggest that familiarity with a distracting talker can be beneficial, but that a high degree of familiarity with the talker is necessary for this benefit to occur. The results of the present study, by contrast, suggest that familiarity with a distracting talker is not beneficial but rather harmful, and that moderate instead of high talker familiarity is necessary for this familiarity effect to occur.

Notably, the present study used an irrelevant-speech task to investigate the distraction by familiar and unfamiliar task-irrelevant talkers. To succeed in the irrelevant-speech task, selective attention to items in working memory is needed (for a review on the interaction of attention and working memory, see Awh et al., 2006). The previous studies, by contrast, used concurrent-speech tasks. For example, Johnsrude et al. (2013) presented listeners with two concurrent sentences (from the Coordinate Response Measure corpus; Bolia et al., 2000) of the form “Ready [call sign], go to [color] [number] now” (e.g., “Ready Baron, go to green six now“) and asked them to report the color and number word from the target sentence. This task creates much lower working-memory load than the irrelevant-speech task used in the present study in which listeners had to keep the serial order of nine digits in memory while ignoring a task-irrelevant sentence. Our results therefore suggest that, while talker familiarity reduces the distraction by irrelevant concurrent speech, it increases the disruption of working memory for target speech.

### Perceptual filtering versus attentional capture of familiar distractors

In the literature, two opposing effects of familiarity with a distracting stimulus have been described. One line of research suggests that advance knowledge about the distractor can enable listeners to form an efficient perceptual filter (e.g., Röer et al., 2015). To suppress distraction by familiar stimuli, the filter can be applied even before the distractor appears (e.g., Noonan et al., 2016; Ruff and Driver, 2006). However, such proactive suppression of the distractor has been found to produce behavioral costs when the distractor is expected but not presented (Marini et al., 2013). Our results clearly speak against proactive filtering in the case of familiar voices because we observed stronger working-memory disruption by a (moderately) familiar versus unfamiliar distracting talker and no modulation of this effect by whether or not a distracting sentence was presented.

Another line of research suggests that familiarity with a distracting stimulus is not beneficial but rather costly for working memory (e.g., Ellermeier et al., 2015; Röer et al., 2013). This is because familiar distractors can automatically capture attention resources and draw them away from items in working memory (Cowan, 1998). While the attentional capture theory predicts stronger working-memory disruption by familiar than unfamiliar talkers, it cannot explain why we only observed a familiarity effect in the student group. If anything, the familiar talker should be more salient for family members and friends who should therefore be more susceptible to attentional capture by the familiar talker’s task-irrelevant speech (but see Gaspelin and Luck, 2019 for a recent review on the suppression of salient stimuli). Furthermore, the attentional capture theory is difficult to reconcile with our finding that the mere expectation of a moderately familiar distractor enhances memory disruption.

### Uncertainty about vocal identity causes working-memory disruption

Our results rather speak to more general differences in how familiar and unfamiliar talkers are processed (for a recent review, see Maguinness et al., 2018). We argue that the disparity of findings both within and across studies can be explained by a model that takes these differences into account, in particularly with regard to how familiarity shapes the representation of vocal identity.

Figure 4 summarizes and illustrates these differences in the representation of familiar and unfamiliar voices, and how these differences may relate to working-memory disruption and distraction by concurrent speech. Consistent with recent advances in voice-identity research (Lavan et al., 2018a), we argue that a high degree of talker familiarity is needed to arrive at a stable representation of vocal identity. Moderate talker familiarity, however, creates uncertainty about vocal identity which, in turn, causes disruption of working memory and distraction by concurrent speech.

**Figure 4.**
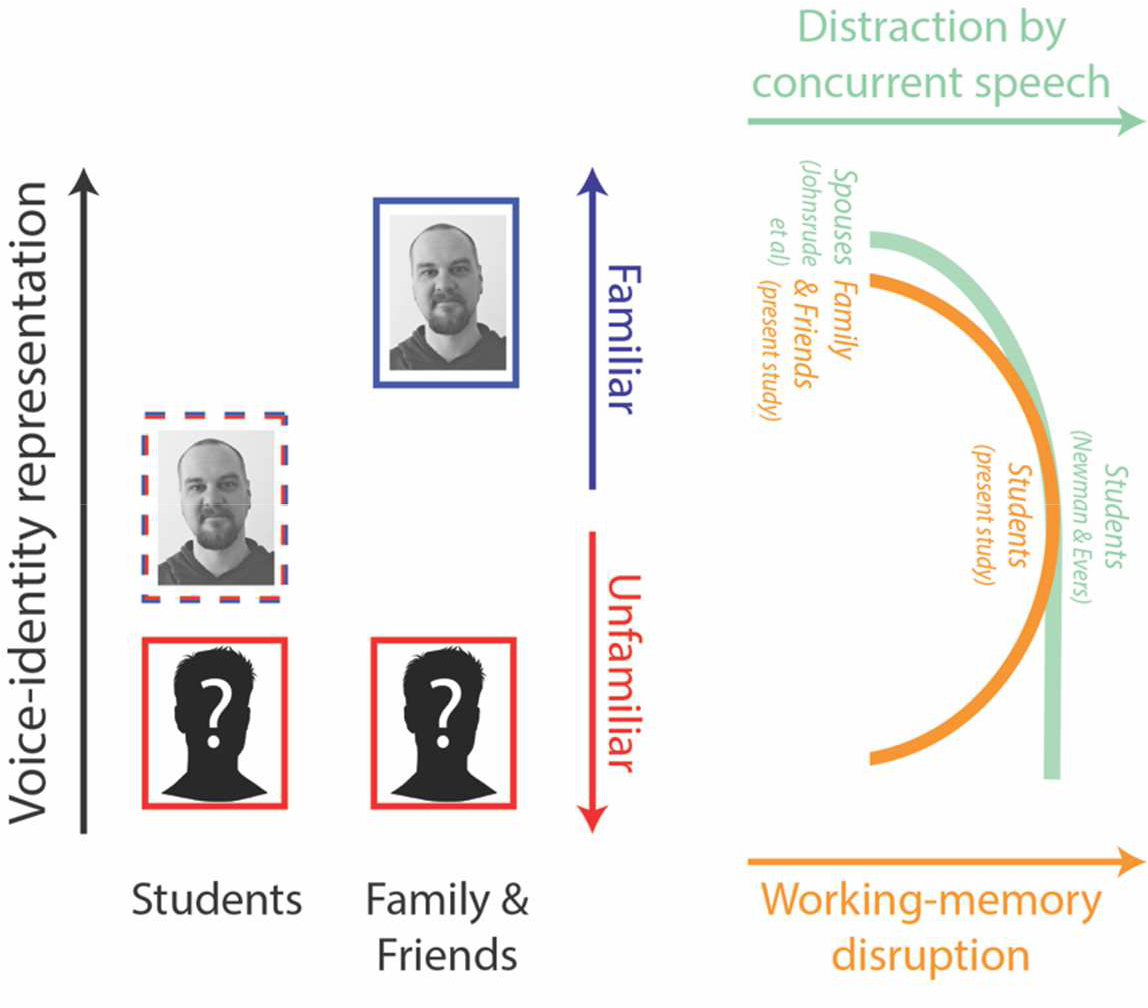
A model of the relationship between talker familiarity and voice-identity representation. Critically, the model assumes insufficient voice-identity representation for moderately familiar talkers causing uncertainty about their vocal identity. This model is capable of explaining the seemingly disparate effects that the degree of familiarity with a to-be-ignored talker has on the disruption of working memory (present study) and the distraction by concurrent speech (Johnsrude et al., 2013; Newman and Evers, 2007). See text for details.

Both the present and previous findings (Johnsrude et al., 2013; Newman and Evers, 2007) have shown that the effect of talker familiarity is not based on a simple dichotomy of familiar versus unfamiliar voices, but that it is rather the degree of familiarity that determines the effect of talker familiarity. We argue that students experienced stronger working-memory disruption by the familiar talker not only because of their limited amount of exposure to the talker’s voice, but, critically, because voice exposure was limited to a very specific context (i.e., classroom teaching). Family members and friends, by contrast, have heard the familiar talker’s voice in a much wider range of contexts. Recent work (Lavan et al., 2018a; 2018b) suggests that it is this variance in previous voice encounters that enables listeners to form a stable percept of person identity.

Of note, talker familiarity was not made explicit in the present study. That is, listeners were not told that they would hear a familiar voice. It is reasonable to assume that family members and friends nevertheless recognized the familiar talker, whereas students remained uncertain about the familiar talker’s identity. Resolving uncertainty about a distractor has been shown to be cognitively effortful (e.g., Geyer et al., 2006; Kerzel and Barras, 2016). In the case of unfamiliar distracting talkers, uncertainty was likely minimal since listeners had little to no expectation about their vocal identity. The difference in vocal uncertainty can thus explain why students experienced stronger working-memory disruption by a moderately familiar than an unfamiliar distracting talker. Critically, this explanation also holds for our finding that the mere expectation of a moderately familiar distractor can cause working-memory disruption.

In the case of concurrent speech, listeners are similarly distracted by an unfamiliar and a moderately familiar talker (Newman and Evers, 2007). Possibly, this is because listeners need more and more variable experience with a talker’s voice to arrive at a representation of vocal identity that is sufficiently reliable to alleviate distraction by irrelevant concurrent speech. This explanation is consistent with a recent extension (Maguinness et al., 2018) of the prototype model of voice-identity processing (Lavner et al., 2001). While listeners can recognize familiar talkers based on stored reference patterns of their vocal identities, such reference patterns need to be yet established for unfamiliar talkers and may not suffice robust identity recognition of moderately familiar talkers. Hence, for both unfamiliar and moderately familiar talkers, additional voice exposure is required.

Several studies have shown a link between voice-identity recognition and speech comprehension (Levi et al., 2011; Magnuson et al., 1995; Nygaard et al., 1994; Nygaard and Pisoni, 1998; but see Holmes et al, 2018): listeners are better at comprehending target speech when they had previously learned to identify the target talker by their voice. It is likely that voice-identity information also helps listeners to attenuate distraction by irrelevant concurrent speech. The findings by Newman and Evers (2007), however, suggest that it is not sufficient to make the identity of the distracting talker explicit. It rather seems that extensive experience with a talker’s voice is needed to overcome immature representation of vocal identity and benefit from familiarity with a distracting talker.

## Conclusions

Here, we demonstrated that moderate, but not high, familiarity with a distracting talker deteriorates working memory of target speech. We propose a model that can explain both the present and previous findings on the distraction by talker familiarity by taking into account how voice familiarity shapes the representation of vocal identity.

## Acknowledgments

We would like to thank all students, family members, and friends who have participated in this experiment, as well as two anonymous reviewers for their valuable comments on an earlier version of the manuscript.

